# Preliminary investigation of Incorporation and Viability of Crystal Violet in *Aspergillus flavus*

**DOI:** 10.1101/091108

**Authors:** Luanda Caroline Parreira de Paula, Daniel Guerra Franco, Clarice Rossatto Marchetti, Giovana Cristina Giannesi, Fabiana Fonseca Zanoelo, Carla Santos de Oliveira

## Abstract

The abusive use of antimicrobials have caused resistance in the microorganisms. Therefore, inovations in research of modern microbiology developed the photodynamic inactivation that is non-toxic and do not promote microbian resistance since it has multiples action sites. When a photoactive molecule is activated by light, it becomes more toxic for the microorganism than for the human cells, increasing the phototoxic effect against such cells. We tested the incorporation of the photosensitizer crystal violet by *Aspergillus flavus* and its viability in different periods of time of incubation (6 and 18 hours), concentrations of crystal violet (4μM, 10μM e 15 μM) and different conditions (resting and agitation). We also tested the photoinactivation of *Aspergillus flavus* containing crystal violet. The incorporation rate does not depend on the concentration of CV neither on the condition of incubation. It depends exclusively on the medium, since it presented higher values in lack ingredients medium. The assays of viability of fungi after exposure to crystal violet show that there is fungic growth when samples are maintained in the dark. However, when the culture were submitted to excitation via laser, there was a decrease in the biomass growth, which indicates that there was photodynamic inactivation of the fungi.

## Introduction

Antibiotics have beend used in the treatment of infecious diseases since 1940 when a group of scientists from Oxford University had success with penicilin in the first clinical test. However, the abusive use have promoted resistance of the microrganisms to the antibiotics in the last decades, and it’s considered one of today’s biggest problems (ALMEIDA *et al*., 2006; BENVINDO *et al*., 2008; DUNBAR *et al*., 2008). Nowadays, inovations in the modern microbiology researches, have developed the photodynamic inhibition, a more efficient therapy, non-invasive and non-toxic that do not promote microbian resistance (PARASCA, 2009). Once the treatment is realized in sick tissues, and on it’s healthy borders, there is a minimum possibility of damage in the neighbor healthy cells. Some researches have shown that photoactive molecules when activated by light, are more toxic for the microorganism than for the human cells, increasing the phototoxic effect against such cells (LUKSIENE, 2005). The variety of affected target sites during photoinactivation prevents tolerant strains to prevail, the biggest vantage of the technic compared to traditional antimicrobian chemotherapy (Komerick & Wilson, 2002).

The kind of light is also essential for success in this therapy and this must be as close to the maximum of absorption of the photosensitizer (TARDIVO, 2005). Photodynamic therapy uses laser as light beam. It’s first use was with argonium of X488 nm and X515 nm, by Mester *et al*. (PINHEIRO, 1992). The laser has characteristics that distinguish from the natural light: the coerence, that means all the waves in the same length, monochromaticity, a pure light with only one color and unidirectional, which possess one direction (PINHEIRO, 2010).

The photosensitizer crystal violet was chosen for this research (Figure 1). It belongs to the family of the triarylmethanes with positive charge in physiological pH, and has photophysical and photochemical characteristics investigated for the application in photodynamic therapy in tumors. Most part of the photosensitizers studied for cancer treatment or others microbians agents are composed of porphyrins, phenothiazines, acridines, phthalocyanines, chlorins, protoporphyrin, haematoporphyrin among others (ENGELMAM, 2005).

**Figure 1:**
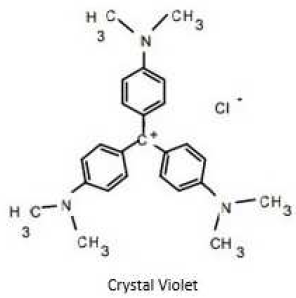
Molecular structure of crystal violet

The photosensitizers must have specific characteristics, such as having an absortion of light in the necessary intensity and wave-length, being efficient in the formation of singlet oxygen, having a low toxicity in the dark, fast elimination by the organism and having specificity for suitable cells sites.

The photodynamic therapy started in the XX century with studies of photosensitizers in paramecium. In another study with acridine, an effective substance against malaria, it was observed that the use of this in culture médium with a certain concentration when exposed to light, the organism obtained toxicity. Then, they analysed that the light and acridine together increased the toxicity rate on paramecium (PINHEIRO, 2010.

It’s a two step therapy. First, there is the accumulation of the photosensitizer agent in the target cells, that are infected cells, this accumulation can be systemic, oral or intravenous. Secondly, the agent sensibilized cell receives a light source with a wavelength that matches the absortion spectrum of the photosensitizer.

When the photon excites the photosensitizer, it is elevated to a more energized state, forming different molecules that are more oxydative or reductive. Once oxygen goes from it’s fundamental state to reactive state, it’s called singlet oxygen. Therefore, when oxygen is excited it reacts with abundant electrons elements, like mitochondria, lysosomes, guanine from DNA, causing oxydation of macromolecules like lipid, aminoacids, which promotes necrosis and tissue death (KESSEL, 2004). There is also a liberation of mediators through immune response unleashed by the sensibilization of the infected tissue. These responses cause destruction of organelles’ membranes, change of selective permeability of the cell and composition of intracellular medium.

There are two types of photosensitizer’s action mechanisms. In the type I the photosensitizer reacts directly in the substrate, which will transmit the electrons to the molecular oxygen. For type II, firstly there is a transferation of electron to molecular oxygen, forming then the free radicals that will oxidize the substrate. These action mechanisms are oxygen, substrate and photosensitizer dependents (GIROTTI, 2001).

In this study we tested the incorporation of the photosensitizer crystal violet by *Aspergillus flavus* and its viability. The incorporation and viability were tested in different times of incubation (6 and 18 hours), concentration (4μM, 10μM e 15 μM), in rest and under agitation conditions. We also tested the photoinactivation of *Aspergillus flavus*containing crystal violet

## Matherials and Methods

### Cultivation and mantaining of fungi

We used an isolated from *Aspergillus flavus*, which is found stored in Micoteca-UFMS. From the pure isolated, we did the inoculum in tubes. The total pureness of the colony is obtained through the cultivation generated from an unique conidia and named “monosporic culture”. Forthe obtaining of this monosporic culture, we collected the suspension of conidia from the pure culture for seeding in Petri dishes containing PDA medium (potato, dextrose and agar) and incubated at 30°C.

This is an important and simple procedure to assure the pureness or the virulelnce from the isolated (SANTOS *et al*. 2009). We prepared microscope slides from the monosporic culture for microculture and from microscope analysis the fungi was characterized with support of the identification key of fungi Illustrated Genere of Imperfect Fungi, 3° ed., Barnett 1972.

Procedures: *Aspergillus flavus* was inoculated in assay tubes containing inclinated solid culture medium (PDA). After growth of fungi, about five days, we prepared a suspension of conidia adding 15mL of destilated water in the culture, rubbing with the handle. From the conidia suspension, we realized the incorporation test (described below) with the inoculum of 1mL of suspension in each one of the Erlenmeyers containing the samples.

### Incorporation of Crystal Violet

For the incorporation of crystal violet, we used two kinds of liquid culture medium for growth of fungi as described on the table 1 below:.

**Table 1.**
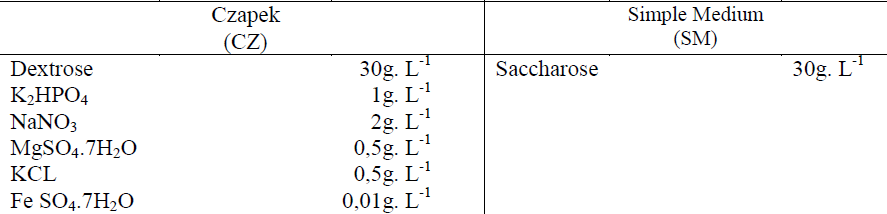
Compounds used as source of nutrients fo preparation of two kinds of liquid culture medium, Czapek medium (CZ) and simple medium (SM).

Medium Preparation: The liquid medium was prepared in Erlenmeyers, wich were autoclaved and after cooled, the assay was initiated. The assays were made in triplicate and two kinds of solutions were prepared: The control, containing the liquid medium plus photosensitizer without the fungi, and in the samples both fungi, photosensitizer and medium were mixed.

In each test we used a concentration of crystal violet, mediated for a period of incubation and condition for fungi, then, we added 1mL of suspension of conidia and incubated at 30°C under rest and agitation conditions.

The incorporation was tested in various assays, ranging time (6h and 18h), conditions (resting and agitation) and concentration of violet crystal (4μM, 10 μM and 15 μM). Then, we took 2 mL of solution from Erlenmeyer and transferred to eppendorfs for centrifugation for 10 minutes and 1000 rpm (Centrifuge 5418).

After this process, the supernatant was taken to ELISA plates or cuvettes for reading in spectrophotometer microplate reader Max 384 plus or Thermo-Scientific spectrophotometer, Genesys 10S UV-VIS, respectively, in 500nm and 580nm wavelength.

The formula used for obtaining of incorporation rate in percentage of photosensitizer absorbed by fungi is represented below:

Incorporation rate = 100-(sample absorption*100/control absorption)

Using this formula, we calculated the averages and generated the graphics for sampling values.

### Viability of Crystal Violet

#### Liquid medium and incorporation solution’s preparation

We prepared liquid medium for the viability test (Table 1), and took it to Erlenmeyers properly autoclaved. After cooling we started the assay.

The assay was made in triplicate and two kinds of solution were prepared: control (fungi in the liquid medium with no photosensitizer) and sample (fungi in the liquid medium plus photosensitizer). In the samples we added the photosensitizer violet crystal and 1mL of suspension of monosporic culture. We tested the viability through 6h and 18h, different concentrations of crystal violet (4μM, 10μM and 15μM), under resting and agitation conditions at 30°C.

#### Solid Medium’s Preparation

It was necessary to prepare solid medium for the viability test to analyse the possible growth of fungi. The solid medium has the same composition of liquid medium with the addition of agar in the solution (Table 2):

**Table 2:** Ingredients for preparation of solid medium

| Czapek<br>(CZ) |  | Simple Medium<br>(SM) |  |
| --- | --- | --- | --- |
| Dextrose | 30g. L<sup>-1</sup> | Saccharose | 30g. L<sup>-1</sup> |
| K<sub>2</sub>HPO<sub>4</sub> | 1g. L<sup>-1</sup> | Agar | 16g. L<sup>-1</sup> |
| NaNO<sub>3</sub> | 2g. L<sup>-1</sup> |  |  |
| MgSO<sub>4</sub>.7H<sub>2</sub>O | 0,5g. L<sup>-1</sup> |  |  |
| KCL | 0,5g. L<sup>-1</sup> |  |  |
| Fe SO<sub>4</sub>.7H<sub>2</sub>O | 0,01g. L<sup>-1</sup> |  |  |
| Agar | 16g. L<sup>-1</sup> |  |  |

All the material was properly autoclaved and posteriorly in a laminar flow ca`binet, we added 20 mL of medium in each Petri dish. After cooling, the dishes were sealed to avoid contamination.

#### Preparation of Viability Test in the Dark

After the incubation period in liquid medium, we transferred 2mL of the solutions of control and sample for each Petri dishes containing the solid medium autoclaved all inside a laminar flow cabinet. This test was also realized in triplicates. Then, the dishes were sealed again and incubated at 30°C in periods of 5 to 7 days.

#### Test of Viability’ Preparation after Luminous Excitation

When the period of incubation in liquid medium was finished, we transferred 2 mL of each incubation solution of both control and sample solutions to Petri dishes containing solid medium properly autoclaved. We made this process in a laminar flow cabin*et al*so in triplicate.

The dishes containing solid medium and solution of spores were maintained at 30°C. During this time the solid medium absorbs both control and sample solutions added. Next, the dishes were unsaddle and washed with 3mL of sodium phosphate Table 2: Ingredients for preparation of solid medium. buffer solution, to eliminate possible excess of solutions. Then, only the sample solutions were submitted to a laser beam for 10 minutes with intervals of 1 minute.

The distance between the dish and laser was about 1cm. After this, the dishes were sealed again and the growth of fungi were analysed during 5 to 7 days at 30°C. The control solutions were also removed from the incubator and washed with the same buffer and then replaced in the incubator. However, it was not submitted to the laser beam.

### Biomass

#### Viability

We accomplished tests in triplicate with concentrations of 4μM, 10 μM and 15 μM of violet crystal. The fungic biomass was obtained with the weighing of the Petri dishes with medium (Table 2) both before and after the incubation assay in the absence and presence of light watching the growth of fungi. This weighing was accomplished in intervals of 7 days through a period of 21 days. From the mass values obtained from fungi, we determined the respective average for each concentration of crystal violet and it’s ranges of fungi’s growth.

#### Dry Mass

To calculate the dry mass we made a ordinary laboratory procedure, wich is used to know the real value of the biomass in one organism, disregarding the existent content of water. It is a long procedure, because the permanence inside heater with adequate temperature that leads to a slow dehydration of mycelium. Generally, we measure biomass until the values become constant, then we have the dry mass value.

We realizated the incorporation of photosensitizer by fungi in liquid medium and transferred 2mL of this culture for the solid medium. When the growth was stabilized, in about 5 days, it was made the rasping of fungi removing the hyphae from the substrate (biomass). The biomass was deposited in to Petri dishes previously weighed. Then, it was transferred to heaters at temperatures of 45°C and 50°C, and daily the dishes were heighed until stability of weight. The stability is given by the constancy of weight that in this case was of three days of measurement.

When constant values were reached, we disregarded the weigth of the dish to obtain the values of biomass. The experiments were realized in triplicate.

### Results and discussion

#### Incorporation of Crystal Violet

The incorporation of photosensitizer by fungi was tested in different concentrations, (4μM, 10 μM and 15 μM), mediums (CZ and SM), time intervals (6 hours and 8 hours) and conditions (resting and agitation). The results of absorption in a concentration of 4μM were significantly good. When the fungi was submitted to agitation conditions, for 18 hours in SM medium, the average absorption was 93%. The second best result in a 4μM concentration was during 6 hours, resting in SM (mean ± SD = 82% ± 2.65%), followed by 18 hours of growth, under agitation, in CZ medium (mean ± SD = 75% ± 7.09%) All the absorption results in 4μM concentration of photosensitizer are shown in figure 2.

**Figure 2.**
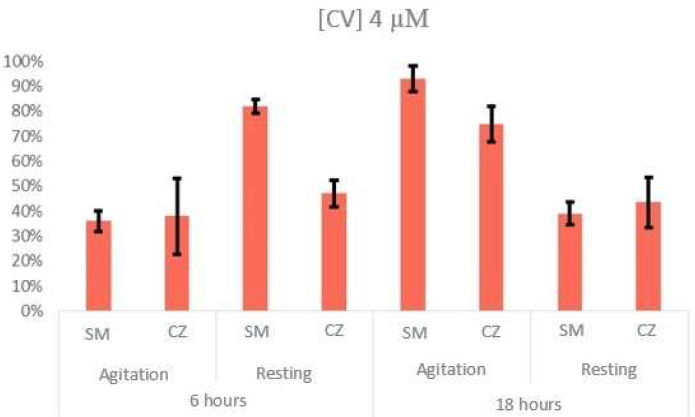
Absorption rate of 4μM of photosensitizer by fungi in different times, conditions and mediums.

On the other hand, the use of photosensitizer in a 10μM concentration showed the best results of absorption. When the growth was made through 18 hours, under agitration using SM medium, there was an average absorption of 93%, and in the same time interval and condition but in CZ medium, the fungi absorbed an average of 74% of the photosensitizer. When submitted to 18 hours of growth under resting condition, there was similar results in both SM and CZ medium (mean ± SD = 82% ± 5.69% and mean ± SD = 80% ± 6.81%, respectively). The results of absorption in a 10 concentration are shown in the figure 3.

**Figure 3:**
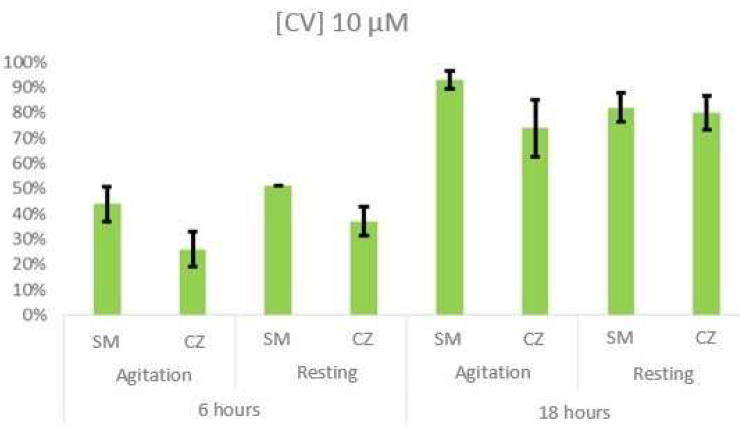
Absorption rate of 10μM of photosensitizer by fungi in different times, conditions and mediums.

There was clearly a considering absorption of the photosensitizer in the data presented so far. The averages of the absorption in a 15μM concentration of photosensitizer were significantly lower than others. Under conditions of agitation through 6 hours of growth, *A. flavus* absorbed 39% in SM and 13% in CZ, and 43.83% under resting in SM medium. At last, through 18 hours of growth, under resting in SM medium, the filamentous fungi absorbed only 30.03% of photosensitizer. The figure 4 below representes the data in a 15μM concentration of crystal violet.

**Figure 4:**
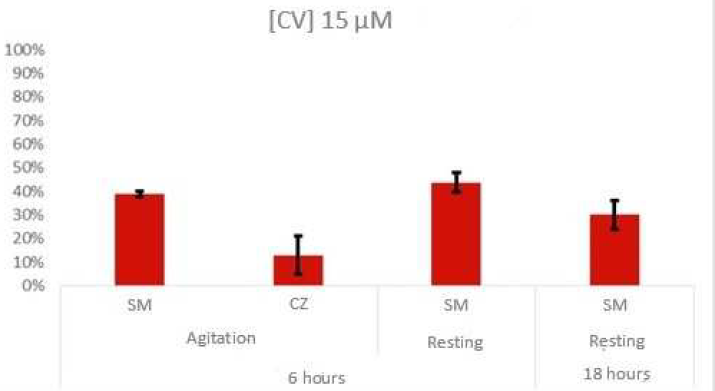
Absorption rate of 15μM of photosensitizer by fungi in different times, conditions and mediums.

The best results of absorption analysed were using 4μM and 10μM under agitation conditions, through 18 hours in SM. Both presented 93% of absorption of the crystal violet. Obviously 93% of 10μM is higher than 93% of 4μM, therefore, using 10μM of photosensitizer crystal violet makes fungi absorb a higher quantity. It is possible to observe that an increase in time makes absorption of CV by fungi better. We have seen that *A. flavus* absorbed a higher rate of crystal violet in SM. A hypothesis suggests that due to the absence of salt in SM, the fungi is induced to search for any kind of salt in the medium, finding only CV that is a cationic salt. It absorbed crystal violet as a source of food in simple medium.

#### Viability of Crystal Violet

As observed, *A. flavus* has a high of absorption rate of the photosensitizer. Therefore, we looked after to know if crystal violet could somehow cause injuries to the mycelium. We realized tests of viability in different variations of time, mediums and conditions, as described in methodology. There was incubations with period of 6 hours under resting and 18 hours under resting and agitation.

When fungi was inoculated from liquid simple medium to solid simple medium, there was no growth of mycelium in both control and sample, as shown in figure 5.

**Figure 5:**
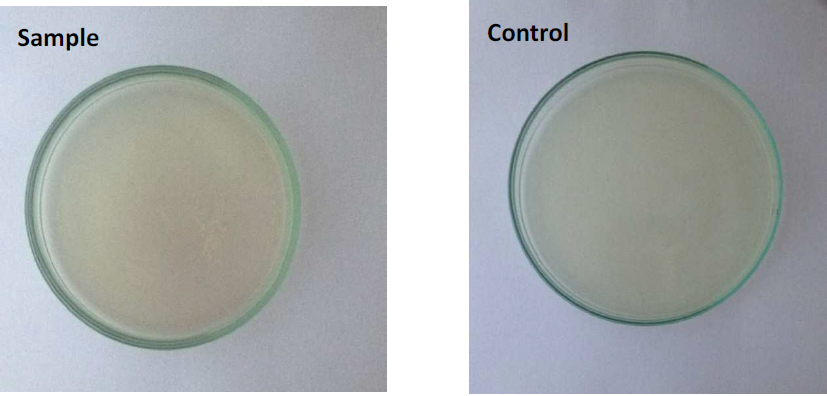
Incubation of *A. ñavus* in liquid simple medium and posteriorly growth of culture in solid simple medium with incubation of [CV] = 4μM.

From these results, we elaborated only CZ solid medium for tests of viability. Besides liquid simple medium is good for crystal violet absorption, when transferred to solid simple medium, fungi could not resist the abscence of nutrients for so long. For solid CZ medium, we incubated fungi in 6h under resting and 18 hours under resting and agitation in simple medium When inoculated in solid CZ medium, fungi showed growth in all triplicates of all concentrations of CV in both control and samples as shown in figure 6. Therefore, we analysed the dry mass of mycelium as described in methodology to determine if the photosensitizer had any effects to mycelium. The results of dry mass in all three concentrations are shown in Table 3 below.

**Table 3:** Procedure of dry mass for assay in 18 hours under agitation, for verification of growth of fungi in samples and controls.

| Biomass - Drymass (g) |  |  |  |
| --- | --- | --- | --- |
| | [CV] = 4μM | [CV] = 10μM | [CV] = 15μM |
| **Sample** | 0,096 | 0,057 | 0,083 |
| **Control** | 0,099 | 0,068 | 0,085 |

**Figure 6:**
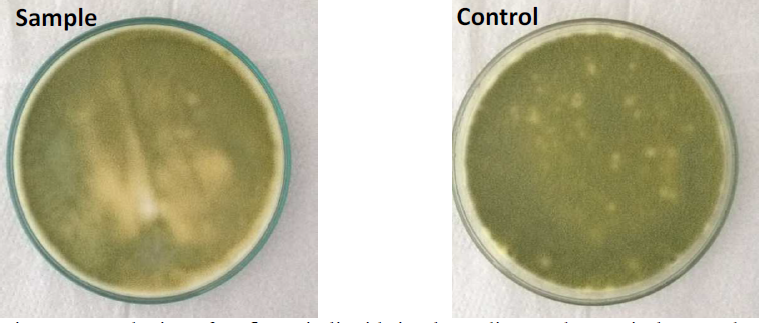
Incubation of *A. ñavus*in liquid simple medium and posteriorly growth of culture in solid CZ medium in Petri dishes with incubation of [CV] = 15μM.

As we can see, there was almost no difference between sample and control in all concentrations of crystal violet. It means that there was no damage to *A. flavus* by crystal violet indeed. As we described before in methodologies, it was realized the crystal violet’s assay in liquid simple medium and posterior inoculum in solid CZ medium after exposure to light with three concentrations of photosensitizer (4μM, 10 μM and 15 μM). To compare the growth of control and sample, we determined the biomass of fungi in intervals of 7 days both in control and samples. After seven days of inoculum, control had grown 52,8%, 57,4% and 51,5% in concentrations of 4μM, 10 μ M and 15 μM respectively. At the fourteenth day, the control’s growth average of all concentrations increased to 78,4 and at the twentieth first day to 83,8%. These results are shown in Table 4.

**Table 4:**
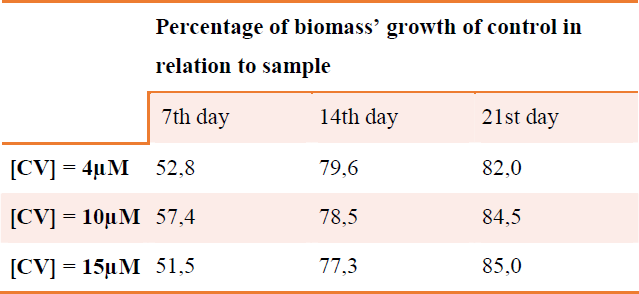
Percentage of values of biomass’ growth of control related to sample.

It is possible to notice in this table an expressive growth of control fungi compared to fungi submitted to excitation via laser. The sample fungi did not progressed with growth, demonstrating that there was photodynamic inactivation of pathogenic fungi *A. flavus.* The studies of viability of fungi after incorporation of CV and maintained in the dark showed that growth was equal both in samples and controls. However, when photosensitizer crystal violet is excited by light irradiation, CV produces a series of radical species wich reacts quickly with molecules around and causes lysis of chemical bonds, formation of highly reactive intermediates, production of toxic molecules and radical redox reactions. (BHASIKUTTAN, A.C *et al*., 1995; CHADHA, R. *et al*., 2013; CHEN, C.C. *et al*, 2006; FAVARO, G. *et al*., 2012; SENTHILKUMAAR, S. *et al*., 2005).

All this perturbation in fungic cell enviroment causes destruction of important biomolecules, promoting reduction of growth’s rate, as observed in the results of viability of fungi after excitation via laser.

## Conclusion

Different conditions for incorporation of photosensitizer by *A. flavus* were detected after 6 and 18 hours of incubation under resting and agitation conditions, in simple medium and CZ medium. A higher exposure to crystal violet promotes a higher absorption of it as the period of incubation increases. There was no significant difference of absorption between resting and agitation conditions. Besides that, simple medium promoted a higher absorption of crystal violet than CZ medium. There was no growth of fungi after inoculum in solid simple medium maintained in the dark. In the assays of viability after excitation via laser, there was growth in all cultures. However, when we analysed the biomass of fungi, there was a higher growth of control fungi in relation to sample fungi, demonstrating that the photodynamic inactivation was promoted. It will be necessary more studies wich search the adequation and efficiency of photosensitizer crystal violet, so it can be used in the treatment of local infections, avoiding use of antimicrobials and the consequent development of resistance.

## Acknowledgments

We acknowledge Rede Pró Centro Oeste – CNPq for the financial support and FUFMS for PIBIC fellowship to LCPP.

## Materials & Correspondence

Prof. Dr. Carla Santos de Oliveira

Laboratório de Bioquímica, CCBS, Universidade Federal de Mato Grosso do Sul Campus Universitário, av. Costa e Silva, s/n, Caixa postal 549, Campo Grande, Mato Grosso do Sul, 79070-900, Brazil.

